## Supplementary File 1 for "Bridging the gap between bioinformatics and the clinical and public health microbiology laboratory: An ISO-accredited genomics workflow for antimicrobial resistance"

### Supplementary Data: Contents

| | |
|---|---|
| **Table S1.** Resistance classes and gene targets determined using synthetic reads | *Page 2* |
| **Table S2.** Steps and logic for *abritAMR* Classification Database updates | *Page 5* |
| **Table S3.** Gene targets and PCR primers for carbapenemase and ESBL detection panel | *Page 8* |
| **Table S4.** Isolates and sequence runs used to determine precision | *Page 9* |
| **Table S5.** Validation results for inferred phenotype in *Salmonella* spp. | *Page 10* |
| **Figure S1.** Reporting logic – AMR gene reporting | *Page 11* |
| **Figure S2.** Reporting logic – Inferred antibiogram reporting | *Page 12* |
| **Figure S3.** Species used in validation of carbapenemase and ESBL detection | *Page 13* |
| **Figure S4.** Gene targets included in validation of carbapenemase and ESBL detection | *Page 14* |
| **Figure S5.** *Enterococcus* and *Staphylococcus* species included in validation of vancomycin resistance gene detection | *Page 15* |
| **Figure S6.** Gene targets included in validation of vancomycin detection | *Page 16* |
| **Figure S7.** Resistance alleles used in validation of allele calling | *Page 17* |
| **Figure S8.** Genera and species included in validation of resistance allele calling | *Page 18* |
| **Figure S9.** Genera and species included in validation of resistance allele calling from synthetic dataset | *Page 19* |
| **Figure S10.** Drug classes with represented in synthetic dataset for the validation of resistance allele calling | *Page 20* |
| **References** | *Page 21* |

#### Table S1. Resistance classes and gene targets determined using synthetic reads

| *AMRFinderPlus* Class | *AMRFinderPlus* Subclass | *abritAMR* Enhanced Subclass | Grouping for reporting | Included in validation panel? |
|---|---|---|---|---|
| Aminoglycoside | By gene: *armA, npmA,* and all *rmt* alleles | Aminoglycoside resistance (ribosomal methyltransferases) | Aminoglycoside resistance (ribosomal methyltransferases) | Y |
| | Amikacin/Gentamicin/ Kanamycin/Tobramycin | Amikacin/Gentamicin/ Kanamycin/Tobramycin | Other aminoglycoside resistance | Y |
| | Amikacin/Kanamycin | Amikacin/Kanamycin | | Y |
| | Amikacin/Kanamycin/Tobramycin | Amikacin/Kanamycin/Tobramycin | | Y |
| | Amikacin/Quinolone | Amikacin/Quinolone | | Y |
| | Amikacin/Tobramycin | Amikacin/Tobramycin | | N |
| | Aminoglycoside | Aminoglycoside | | Y |
| | Gentamicin | Gentamicin | | Y |
| | Gentamicin/Kanamycin/Tobramycin | Gentamicin/Kanamycin/Tobramycin | | Y |
| | Gentamicin/Tobramycin | Gentamicin/Tobramycin | | Y |
| | Hygromycin | Hygromycin | | Y |
| | Kanamycin | Kanamycin | | Y |
| | Kanamycin/Tobramycin | Kanamycin/Tobramycin | | Y |
| | Kasugamycin | Kasugamycin | | N |
| | Spectinomycin | Spectinomycin | | Y |
| | Streptomycin | Streptomycin | | Y |
| | Streptomycin/Spectinomycin | Streptomycin/Spectinomycin | | Y |
| | Tobramycin | Tobramycin | | Y |
| Beta-lactam | Carbapenem | Carbapenemase (MBL)<br>Carbapenemase (OXA-51 family)<br>Carbapenemase (see Appendix 15.1 for classification rules) | Carbapenemase (MBL)<br>Carbapenemase (OXA-51 family)<br>Carbapenemase | Y |

| *AMRFinderPlus* Class | *AMRFinderPlus* Subclass | *abritAMR* Enhanced Subclass | Grouping for reporting | Included in validation panel? |
|---|---|---|---|---|
| | Cephalosporin | ESBL<br>ESBL (AmpC)<br>Beta-lactamase (not carbapenemase or ESBL)<br>(see Appendix 15.1 for classification rules) | ESBL<br>ESBL (AmpC)<br>Beta-lactamase (not carbapenemase or ESBL) | Y |
| | Beta-lactam | Beta-lactamase (not carbapenemase or ESBL)<br>Beta-lactamase (unknown spectrum)<br>Penicillin resistance (*Staphylococcus aureus*) (see Appendix 15.1 for classification rules) | Beta-lactamase (not carbapenemase or ESBL)<br>Beta-lactamase (unknown spectrum)<br>Penicillin resistance (*Staphylococcus aureus*) | Y |
| | Cephalothin | ESBL (AmpC type) | ESBL (AmpC type) | Y |
| | Methicillin | Methicillin | Methicillin | Y |
| Colistin | Colistin | Colistin | Colistin | Y |
| Fosfomycin | Fosfomycin | Fosfomycin | Fosfomycin | Y |
| Fusidic Acid | Fusidic Acid | Fusidic Acid | Fusidic Acid | Y |
| Glycopeptide | Vancomycin | Vancomycin | Vancomycin | Y |
| Lincosamide | Lincosamide | Lincosamide | Macrolide, lincosamide and/or streptogramin resistance | Y |
| Lincosamide/Streptogramin | Lincosamide/Streptogramin | Lincosamide/Streptogramin | Macrolide, lincosamide and/or streptogramin resistance | Y |
| Macrolide | Macrolide | Macrolide | Macrolide, lincosamide and/or streptogramin resistance | Y |
| | Erythromycin | Erythromycin | Macrolide, lincosamide and/or streptogramin resistance | N |
| | Erythromycin/Telithromycin/Tylosin | Erythromycin/Telithromycin/Tylosin | Macrolide, lincosamide and/or streptogramin resistance | N |
| Macrolide/Lincosamide/Streptogramin | Macrolide/Lincosamide/Streptogramin | Macrolide/Lincosamide/Streptogramin | Macrolide, lincosamide and/or streptogramin resistance | Y |
| Macrolide/Pleuromutilin | Lincosamide/Streptogramin/Tiamulin | By gene: *cfr* alleles, *cipA, clbA*: Oxazolidinone & phenicol resistance | Oxazolidinone & phenicol resistance | N |

| *AMRFinderPlus* Class | *AMRFinderPlus* Subclass | *abritAMR* Enhanced Subclass | Grouping for reporting | Included in validation panel? |
|---|---|---|---|---|
| Streptogramin | Streptogramin | Streptogramin | Macrolide, lincosamide and/or streptogramin resistance | N |
| Mupirocin | Mupirocin | Mupirocin | Mupirocin | Y |
| Nitroimidazole | Nitroimidazole | Nitroimidazole | Nitroimidazole | N |
| Phenicol | Chloramphenicol | Chloramphenicol | Chloramphenicol | Y |
| | Chloramphenicol/Florphenicol | Chloramphenicol/Florphenicol | Chloramphenicol/Florphenicol | Y |
| | Phenicol | Phenicol | Phenicol | N |
| Phenicol/ Oxazolidinone | Florfenicol/Oxazolidinone | Oxazolidinone & phenicol resistance (by gene: *optrA*) | Oxazolidinone & phenicol resistance | Y |
| Phenicol/Quinolone | Phenicol/Quinolone | Phenicol/Quinolone | Phenicol/Quinolone | Y |
| Quinolone | Quinolone | Quinolone | Quinolone | Y |
| Rifamycin | Rifampin | Rifampin | Rifampin | Y |
| Sulfonamide | Sulfonamide | Sulfonamide | Sulfonamide | Y |
| Tetracycline | Tetracycline | Tetracycline | Tetracycline | Y |
| Trimethoprim | Trimethoprim | Trimethoprim | Trimethoprim | Y |
| Avilamycin | Avilamycin | Other | Other | Y |
| Bleomycin | Bleomycin, Zorbamycin | Other | Other | Y |
| Pleuromutilin | Tiamulin | Other | Other | N |
| Streptothricin | Streptothricin | Other | Other | Y |
| Tetracenomycin | Tetracenomycin | Other | Other | Y |
| Tuberoactinomycin | Viomycin | Other | Other | N |
| Thiostrepton | Thiostrepton | Other | Other | N |

#### Table S2. Steps and logic for *abritAMR* Classification Database updates

After each new *AMRFinderPlus* release (those including new AMR genes or mutations), the *abritAMR* Classification Database is updated by the curators.

An initial classification step is performed according to the following logic for all new AMR genes added in the database update, as described in Table S2 below. This is supplemented by manual review and curation of critical resistance gene families, and other groups not covered by the update logic. Additionally, any changes in existing genes since the last update are reviewed.

| **Step 1: Classification by alleles** | |
|---|---|
| ***AMRFinderPlus* allele** | ***abritAMR* Classification Database Enhanced Subclass** |
| *rmt*, armA, npmA* | Aminoglycoside resistance (ribosomal methyltransferases) |
| *optrA, cfr*, cipA, clb* alleles | Oxazolidinone & phenicol resistance |
| **Step 2: Classification by NCBI Subclass** | |
| ***AMRFinderPlus* Subclass** | ***abritAMR* Classification Database Enhanced Subclass** |
| Carbapenemase | If "metallo" in description → Carbapenemase (MBL);<br>If "OXA-51 family" or allele=OXA-51 → "Carbapenemase (OXA-51 family);<br>Otherwise "Carbapenemase" |
| Cephalosporin | If description includes "class C" → "ESBL (AmpC)";<br>If description includes "extended-spectrum" and NOT "class C" → "ESBL" |
| Beta-lactam | If description includes "broad-spectrum" or "carbenicillin-hydrolyzing" → classified as "Beta-lactamase (not ESBL or carbapenemase)";<br>If *blaZ** alleles -> "Penicillin resistance (*Staphylococcus aureus*)"<br>Other groups curated curated manually; if no phenotype found, called "Beta-lactamase (unknown spectrum)" |
| Colistin | Colistin |
| Vancomycin | Vancomycin |
| Methicillin | Methicillin |
| Fosfomycin | Fosfomycin |
| Quinolone | Quinolone |
| Tetracycline | Tetracycline |
| Trimethoprim | Trimethoprim |
| Rifamycin | Rifamycin |
| Chloramphenicol | Chloramphenicol |
| Chloramphenicol/Florfenicol | Chloramphenicol/Florfenicol |
| Phenicol | Phenicol |
| Phenicol/Quinolone | Phenicol/Quinolone |
| Fusidic acid | Fusidic acid |

| | |
|---|---|
| Mupirocin | Mupirocin |
| Nitroimidazole | Nitroimidazole |
| Amikacin/Gentamicin/Kanamycin/Tobramycin | Other aminoglycoside resistance (non-RMT) |
| Amikacin/Kanamycin | Other aminoglycoside resistance (non-RMT) |
| Amikacin/Kanamycin/Tobramycin | Other aminoglycoside resistance (non-RMT) |
| Amikacin/Quinolone | Other aminoglycoside resistance (non-RMT) |
| Amikacin/Tobramycin | Other aminoglycoside resistance (non-RMT) |
| Aminoglycosides | Other aminoglycoside resistance (non-RMT) |
| Gentamicin | Other aminoglycoside resistance (non-RMT) |
| Gentamicin/Kanamycin/Tobramycin | Other aminoglycoside resistance (non-RMT) |
| Gentamicin/Tobramcyin | Other aminoglycoside resistance (non-RMT) |
| Kanamycin | Other aminoglycoside resistance (non-RMT) |
| Kanamycin/Tobramycin | Other aminoglycoside resistance (non-RMT) |
| Spectinomycin | Other aminoglycoside resistance (non-RMT) |
| Streptomycin | Other aminoglycoside resistance (non-RMT) |
| Streptomycin/Spectinomycin | Other aminoglycoside resistance (non-RMT) |
| Tobramycin | Other aminoglycoside resistance (non-RMT) |
| Erythromycin | Macrolide, lincosamide & streptogramin resistance |
| Erythromycin/Telithromycin | Macrolide, lincosamide & streptogramin resistance |
| Lincosamides | Macrolide, lincosamide & streptogramin resistance |
| Lincosamides/Streptogramin | Macrolide, lincosamide & streptogramin resistance |
| Macrolides | Macrolide, lincosamide & streptogramin resistance |
| Streptogramin | Macrolide, lincosamide & streptogramin resistance |
| Bleomycin | Other |
| Pleuromutilin | Other |
| Avilamycin | Other |
| Streptothricin | Other |
| Tetracenomycin | Other |
| Thiostrepton | Other |
| Tuberactinomycin | Other |

#### Step 3: Manual review and curation

| ***AMRFinderPlus*** **Subclass** | ***abritAMR*** **Classification Database Enhanced Subclass** |
|---|---|
| Carbapenemase, Cephalosporin, Beta-lactam | Classifications reviewed and curated manually<br>Where phenotype unclear from gene family or description, manual search to determine appropriate subclass:<br>• Review GenBank reference sequence record for associated publications and references<br>• Search CARD for record or publication<br>• Search PubMed and Google Scholar for publications<br><br>When no phenotypic data available (insufficient description, no publications, or metagenomic data), classify as 'Beta-lactamase (unknown spectrum)' |

| Step 4: Review changes to existing genes since last update |
| --- |
| Any genes where the class, subclass or description have changed since the last update are also reviewed to determine the significance, and if any changes are required to the Enhanced Subclass |

*, wildcard (i.e. any character/s); RMT, ribosomal methyltransferase; CARD, Comprehensive Antimicrobial Resistance Database (https://card.mcmaster.ca).

#### Table S3. Gene targets and PCR primers for carbapenemase and ESBL detection and allelic typing by Sanger sequencing

| Gene/Target | Primer | Primer sequence | Product size (bp) | Ref |
|---|---|---|---|---|
| blaKPC | KpcF<br>KpcR | ATG TCA CTG TAT CGC CGT C<br>TTA CTG CCC GTT GAC GCC-3' | 845 | (1) |
| blaOXA-48-like | OXA-48F<br>OXA-48R | TTG GTG GCA TCG ATT ATC GG<br>GAG CAC TTC TTT TGT GAT GGC | 743 | (2) |
| blaIMP | IMP-A<br>IMP-B | GAA GGY GTT TAT GTT CAT AC<br>GTA MGT TTC AAG AGT GAT GC | 587 | (3) |
| blaVIM | VIM2004-A<br>VIM2004-B | GTT TGG TCG CAT ATC GCA AC<br>AAT GCG CAG CAC CAG GAT AG | 382 | |
| blaNDM | NDM-F<br>NDM-R | GGG CAG TCG CTT CCA ACG GT<br>GTA GTG CTC AGT GTC GGC AT | 475 | (4) |
| IMP sequencing<br>5'CS-IMP-B | 5'CS<br>IMP-B | GGC ATC CAA GCA GCA AG<br>GTA MGT TTC AAG AGT GAT GC | 793 | (3) |
| VIM 1 amplification & sequencing | Vim-1F<br>Vim-1R | TTA TGG AGC AGC AAC GAT GT<br>CAA AAG TCC CGC TCC AAC GA | 920 | |
| VIM-2 amplification & sequencing | Vim-2F<br>Vim-2R | AAA GTT ATG CCG CAC TCA CC<br>TGC AAC TTC ATG TTA TGC CG | 865 | (5) |
| VIM-2 seq | Vim-2sF<br>Vim-2sR | TCG ACG GTG ATG CGT ACG TT<br>TTG ATG TCC TTC GGG CGG CT | 865 | |
| NDM seq | NDMLF<br>NDMLR | ATG GAA TTG CCC AAT ATT ATG CAC<br>TCA GCG CAG CTT GTC GGC | 813 | (6) |
| blaMOX | MOXF<br>MOXR | GCT GCT CAA GGA GCA CAG GAT<br>CAC ATT GAC ATA GGT GTG GTG C | 520 | |
| blaCIT | CITF<br>CITR | TGG CCA GAA CTG ACA GGC AAA<br>TTT CTC CTG AAC GTG GCT GGC | 462 | |
| blaDHA | DHAF<br>DHAR | AAC TTT CAC AGG TGT GCT GGG T<br>CCG TAC GCA TAC TGG CTT TGC | 405 | |
| blaACC | ACCF<br>ACCR | AAC AGC CTC AGC AGC CGG TTA<br>TTC GCC GCA ATC ATC CCT AGC | 346 | (7) |
| blaEBC | EBCF<br>EBCR | TCG GTA AAG CCG ATG TTG CGG<br>CTT CCA CTG CGG CTG CCA GTT | 302 | |
| blaFOX | FOXF<br>FOXR | AAC ATG GGG TAT CAG GGA GAT G<br>CAA AGC GCG TAA CCG GAT TGG' | 190 | |
| blaOXA-23-like | OXA-23F<br>OXA-23R | GAT CGG ATT GGA GAA CCA GA<br>ATT TCT GAC CGC ATT TCC AT | 5001 | |
| blaOXA-24-like | OXA-24F<br>OXA-24R | GGT TAG TTG GCC CCC TTA AA<br>AGT TGA GCG AAA AGG GGA TT | 246 | (8) |
| blaOXA-51-like | OXA-51F<br>OXA-51R | TAA TGC TTT GATCGG CCT TG<br>TGG ATT GCA CTT CAT CTT GG | 353 | |
| blaOXA-58-like | OXA-58F<br>OXA-58R | AAG TAT TGG GGC TTG TGC TG<br>CCC CTC TGC GCT CTA CAT AC | 599 | |

#### Table S4. Isolates and sequence runs used to determine precision

| **Species** | **Average genome length Mbp (Range)** | **Average % GC (Range)** | **Gram stain** |
|---|---|---|---|
| *Clostridium difficile* | 4.23 (4.05-4.46) | 28.81 (28.35-29.20) | Positive |
| *Campylobacter jejuni* | 1.67 (1.61-1.85) | 30.41 (30.18-30.70) | Negative |
| *Listeria monocytogenes* | 2.96 (2.78-3.24) | 38.02 (37.86-38.30) | Positive |
| *Legionella pneumophila* | 3.43 (2.68-3.79) | 38.34 (38.10-38.60) | Negative |
| *Streptococcus pyogenes* | 1.84 (1.70-1.95) | 38.51 (38.20-38.70) | Positive |
| *Escherichia coli* | 5.07 (4.48-5.87) | 50.69 (50.29-51.20) | Negative |
| *Neisseria meningitidis* | 2.19 (2.14-2.32) | 51.65 (51.30-51.90) | Negative |
| *Salmonella* Typhimurium | 4.82 (4.76-5.18) | 52.15 (52.08-52.30) | Negative |
| *Neisseria gonorrhoeae* | 2.21 (2.15-2.24) | 52.48 (52.40-52.70) | Negative |
| *Klebsiella pneumoniae* | 5.6 (5.09-6.13) | 57.16 (56.54-58.00) | Negative |
| *Mycobacterium tuberculosis* | 4.41 (4.38-4.44) | 65.60 (65.60-65.60) | (Positive) |
| *Pseudomonas aeruginosa* | 6.62 (6.12-7.50) | 66.24 (65.60-66.70) | Negative |
| *Burkholderia lata* | 7.62 (6.42-8.61) | 66.84 (66.60-67.06) | Negative |

#### Table S5. Validation of inferred phenotype for *Salmonella* spp. by *abritAMR*[a]

| Antimicrobial | Accuracy[b] (%, 95%CI) | Sensitivity (%, 95%CI) | Specificity (%, 95%CI) | PPV (%, 95%CI) | NPV (%, 95%CI) |
|---|---|---|---|---|---|
| **Ampicillin (*n*=863)** | 99.1 (98.2-99.6) | 99.2 (97.8-99.8) | 98.9 (97.5-99.7) | 98.7 (97.1-99.6) | 99.4 (98.1-99.9) |
| **Cefotaxime (*n*=862)** | 99.9 (99.4-100) | 100 (92.0-100) | 99.9 (99.3-100) | 97.8 (88.2-99.9) | 100 (99.5-100) |
| **Meropenem[c] (*n*=789)** | 100 (99.5-100) | 100 (15.8-100) | 100 (99.5-100) | 100 (15.8-100) | 100 (99.5-100) |
| **Gentamicin[d] (*n*=846)** | 99.8 (99.1-100) | 100 (87.2-100) | 99.8 (99.1-100) | 93.1 (77.2-99.2) | 100 (99.5-100) |
| **Kanamycin[d] (*n*=864)** | 100 (99.6-100) | 100 (92.3-100) | 100 (99.6-100) | 100 (92.3-100) | 100 (99.6-100) |
| **Streptomycin[d] (n=716)** | 95.5 (93.7-96.9) | 99.2 (97.0-99.9) | 93.7 (91.1-95.7) | 88.8 (84.5-92.3) | 99.6 (98.4-99.9) |
| **Sulfathiazole (*n*=864)** | 98.8 (97.9-99.4) | 97.9 (95.7-99.2) | 99.4 (98.4-99.9) | 99.1 (97.4-99.8) | 98.7 (97.3-99.5) |
| **Trimethoprim (*n*=864)** | 99.5 (98.8-99.9) | 100 (97.5-100) | 99.4 (98.6-99.8) | 97.4 (93.4-99.3) | 100 (99.5-100) |
| **Trimethoprim-Sulfamethoxazole (*n*=864)** | 99.1 (98.2-99.6) | 96.9 (92.3-99.1) | 99.5 (98.6-99.9) | 96.9 (92.3-99.1) | 99.5 (98.6-99.9) |
| **Tetracycline (*n*=859)** | 98.5 (97.4-99.2) | 99.4 (98.0-99.9) | 97.8 (96.1-98.9) | 97.0 (94.7-98.5) | 99.6 (98.5-100) |
| **Chloramphenicol (*n*=858)** | 99.3 (98.5-99.7) | 100 (96.7-100) | 99.2 (98.3-99.7) | 94.8 (89.0-98.1) | 100 (99.5-100) |
| **Azithromycin (*n*=864)** | 99.2 (98.3-99.7) | 85.3 (68.9-95.0) | 99.8 (99.1-100) | 93.5 (78.6-99.2) | 99.4 (98.6-99.8) |
| **Ciprofloxacin[e] (*n*=864)** | 96.8 (95.4-97.8) | 99.6 (98.5-99.9) | 93.5 (90.6-95.7) | 94.7 (92.3-96.5) | 99.5 (98.1-99.9) |
| **Summary performance (all antimicrobials)(95% CI)** | **98.9 (98.7-99.1)** | **98.9 (98.4-99.3)** | **98.9 (98.7-99.1)** | **96.1 (95.2-96.8)** | **99.7 (99.6-99.8)** |

PPV, positive predictive value; NPV, negative predictive value; CI, confidence interval.
[a] Compared to agar dilution using CLSI methods and breakpoints (CLSI M100 2020). Excludes all 'intermediate' results from AST.
[b] Accuracy determined by number of true positives and negatives divided by total number of results (true positive, genotypic and phenotypic resistance; true negative, no AMR mechanisms detected in phenotypically susceptible isolate).
[c] Note only two carbapenemase-producing isolates were available for inclusion in the validation dataset.
[d] Note that aminoglycosides are not used for patient treatment, but included here for surveillance purposes.
[e] For ciprofloxacin, 'true positive' defined as concordant intermediate or resistant results (phenotype and genotype).

#### Figure S1. Reporting logic – AMR gene reporting

This figure details the logic used for the final step of *abritAMR*, to determine whether each result is placed in to 'Reportable' or 'Non-reportable' fields of the Final AMR gene Report output (for clinical microbiology reports).

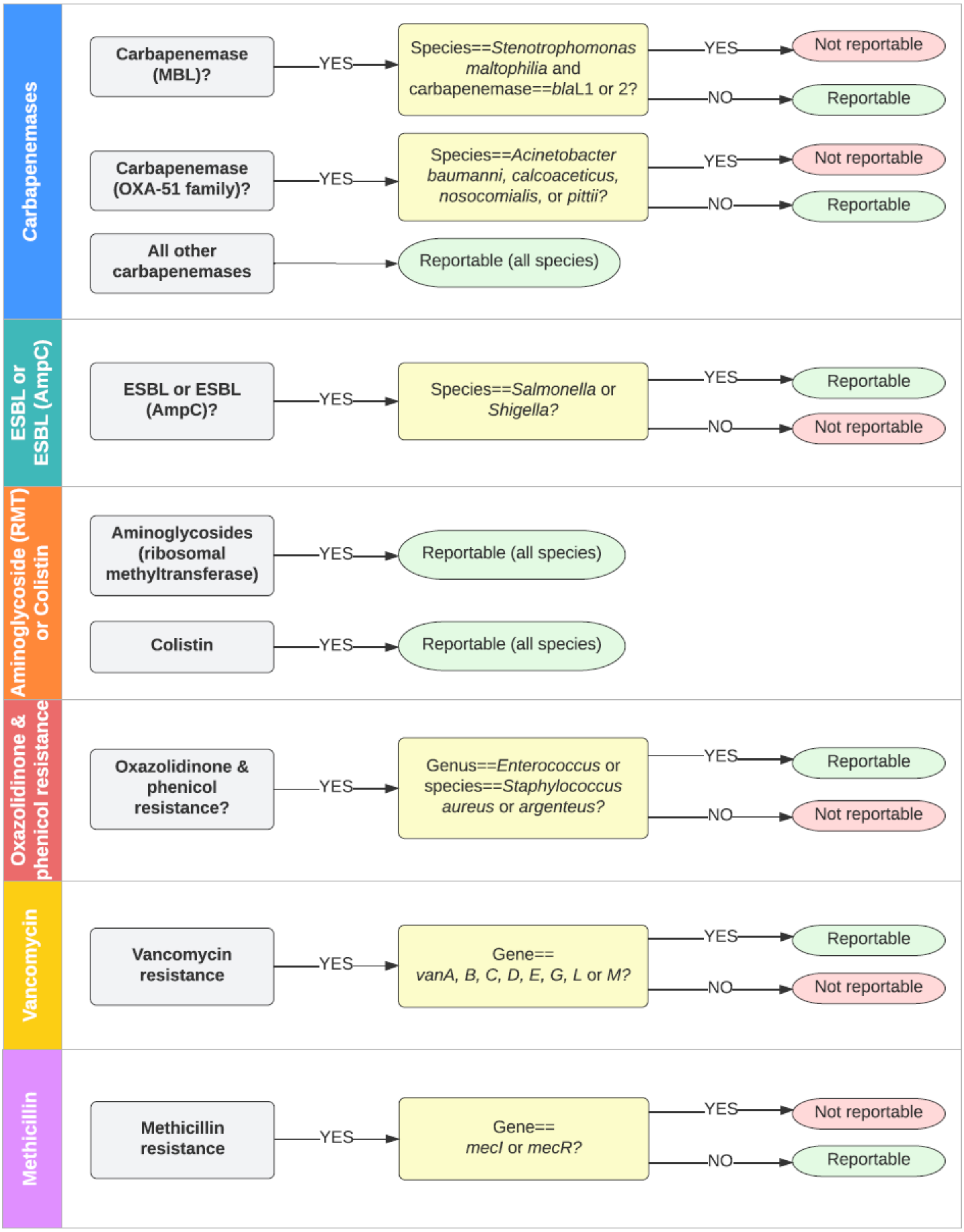

#### Figure S2. Reporting logic – Inferred antibiogram reporting

**IF ANY OF THE FOLLOWING DETECTED >> RESISTANT OTHERWISE SUSCEPTIBLE**

| Antibiotic | Rule |
|---|---|
| Ampicillin | **Any gene from enhanced subclasses** ==Beta-lactamase (narrow spectrum) ==Beta-lactamase (not ESBL or carbapenemase) ==Cefotaxime (ESBL) ==Cefotaxime (AmpC type), or contains "Carbapenem" |
| Cefotaxime (ESBL) | **Any gene from enhanced subclasses** ==ESBL, or contains "Carbapenem" |
| Cefotaxime (AmpC type) | **Any gene from enhanced subclasses** ==ESBL (AmpC type), or contains "Carbapenem" |
| Meropenem | **Any gene from enhanced subclasses** contains "Carbapenemase" except "Carbapenemase (KPC variant)" |
| Tetracycline | **Any gene from enhanced subclass** contains "Tetracycline" |
| Azithromycin | **Any gene or mutation from enhanced subclass:** contains "Azithromycin" |

**IF ANY OF THE FOLLOWING DETECTED>> RESISTANT OTHERWISE SUSCEPTIBLE**

| Antibiotic | Rule |
|---|---|
| Trimethoprim | **Any gene from enhanced subclass** ==Trimethoprim |
| Sulfathiazole | **Any gene from enhanced subclass** ==Sulfonamide |
| Trimethoprim-Sulfamethoxazole | **Any gene from enhanced subclasses** ==Trimethoprim, or ==Sulfonamide |
| Gentamicin | **Any gene from enhanced subclasses** contains "Gentamicin" or contains "Aminoglycoside" |
| Kanamycin | **Any gene from enhanced subclass** contains "Kanamycin" |
| Streptomycin | **Any gene from enhanced subclass** contains "Streptomycin" |
| Chloramphenicol | **Any gene from enhanced subclass** contains "Phenicol" |

**SPECIAL CASES**

**Ciprofloxacin**

**AMR genes or mutations from enhanced subclasses** contain "Ciprofloxacin" or "Quinolone"

No AMR genes or mutations >> **Susceptible**

One AMR gene or mutation >> **Intermediate**

Two or more AMR genes or mutations >>**Resistant**

**REPORTING OF AMR GENE CLASSES ONLY (NO INFERRED ANTIBIOGRAM)**

| | |
|---|---|
| Aminoglycosides (ribosomal methyltransferases) | Reportable mechanism |
| Colistin | Mobile colistin resistance (*mcr*) genes Reportable mechanism |
| Other | AMR genes not reported under any other category; recorded in LIMS, not included in routine reports |

**Figure S3: Species used in validation of carbapenemase and ESBL detection**

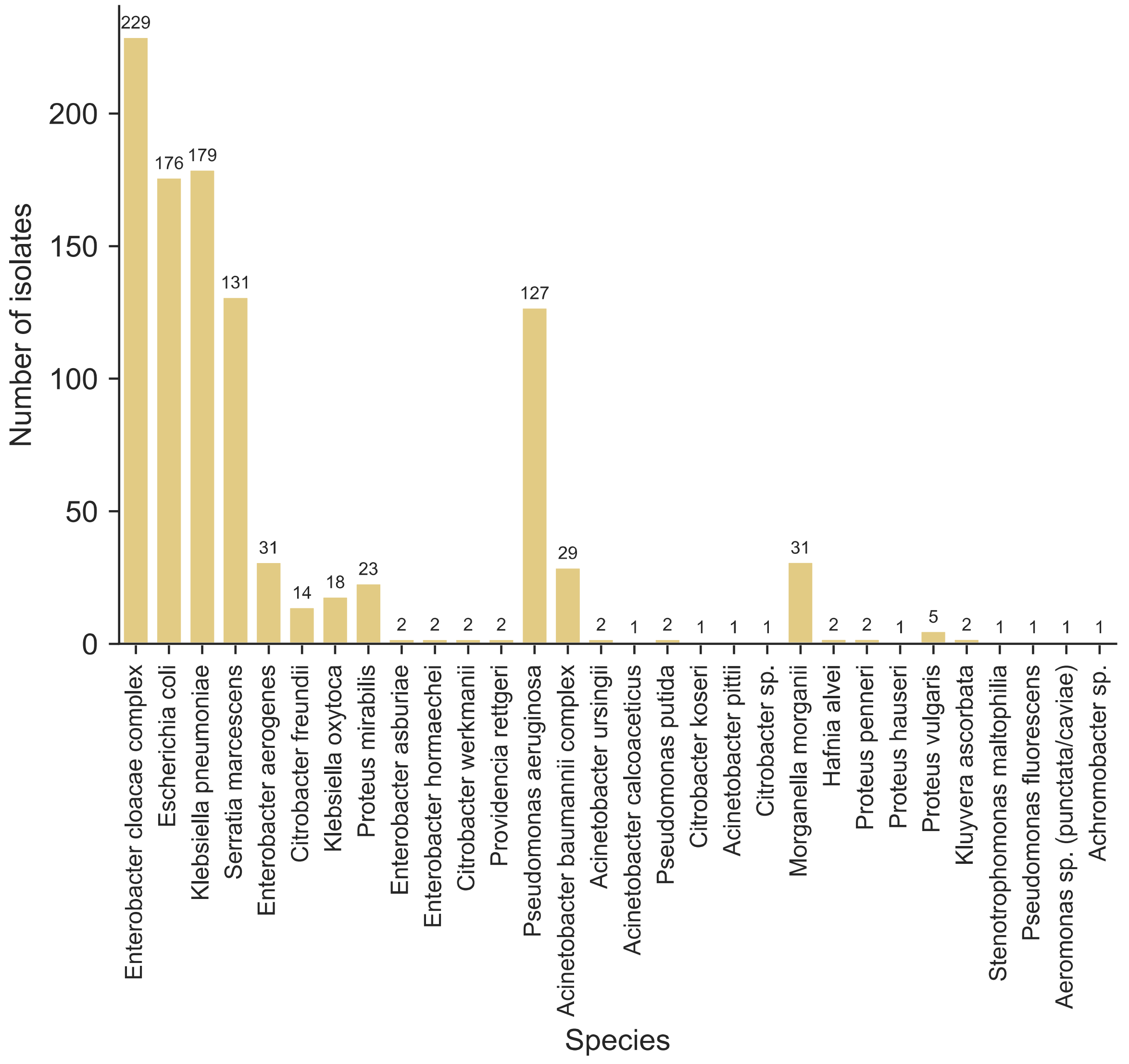

#### Figure S4: Gene targets included in validation of carbapenemase and ESBL detection

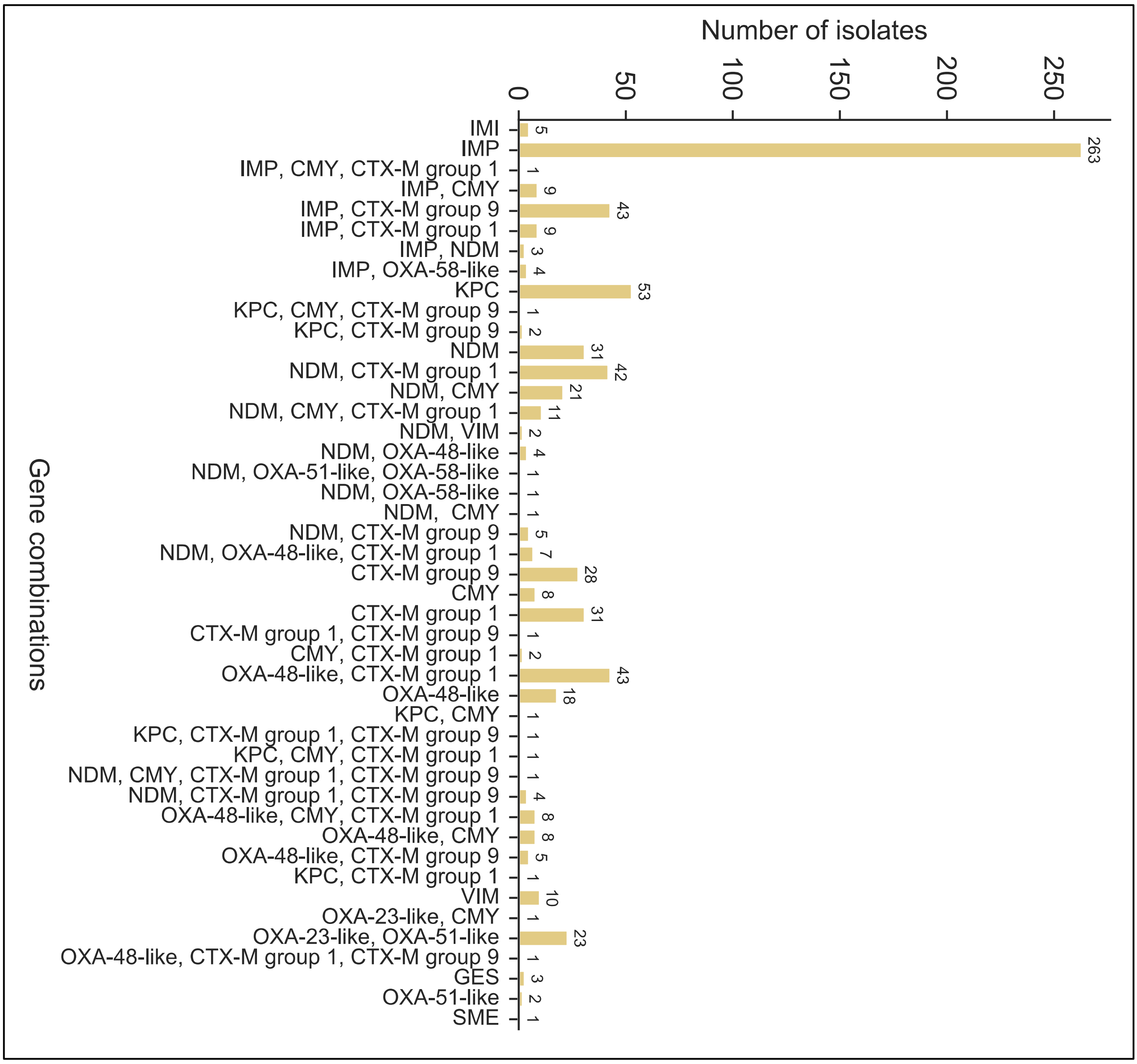

**Figure S5: *Enterococcus* and *Staphylococcus* species included in validation of vancomycin resistance gene detection**

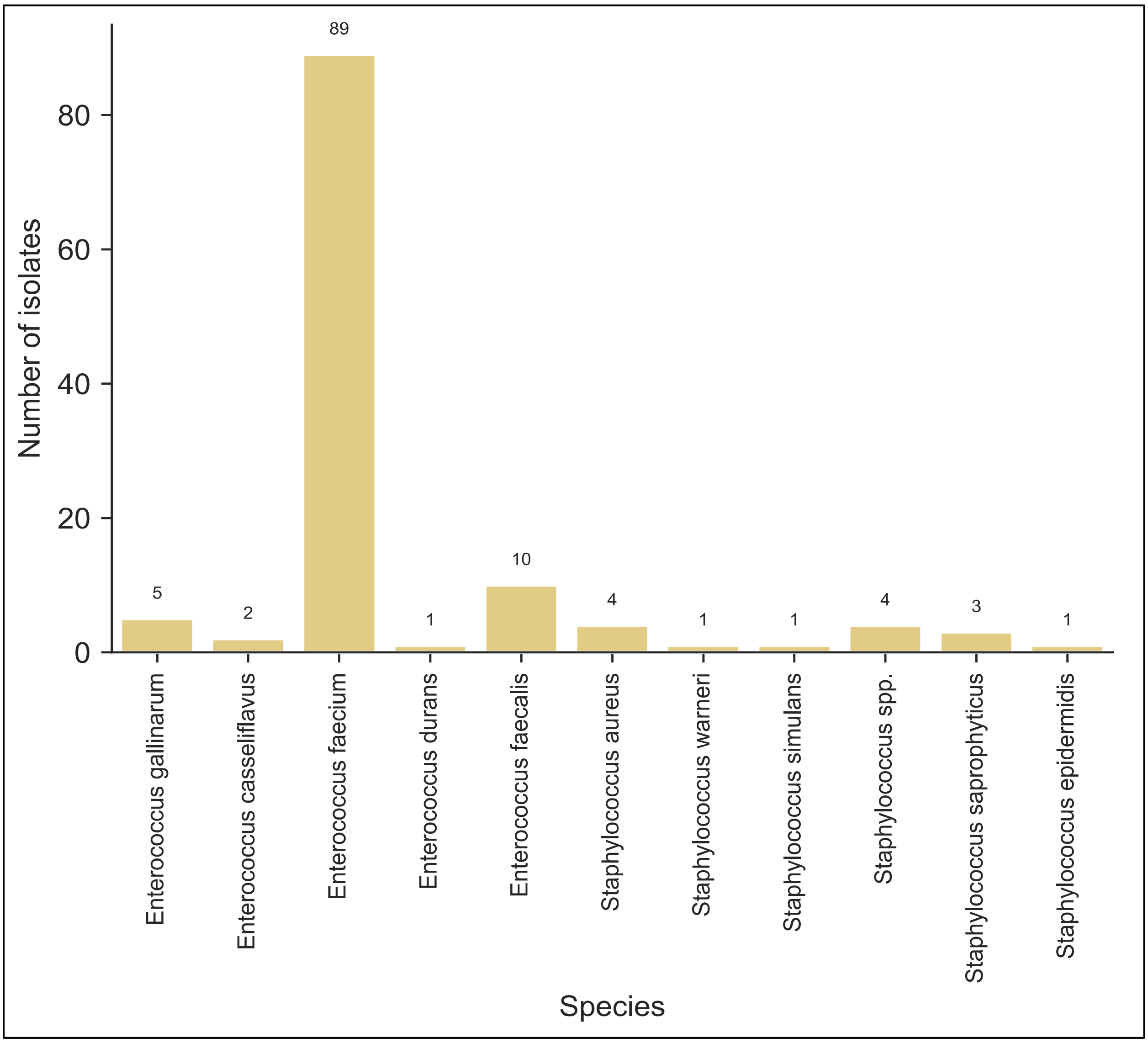

**Figure S6: Gene targets included in validation of vancomycin detection**

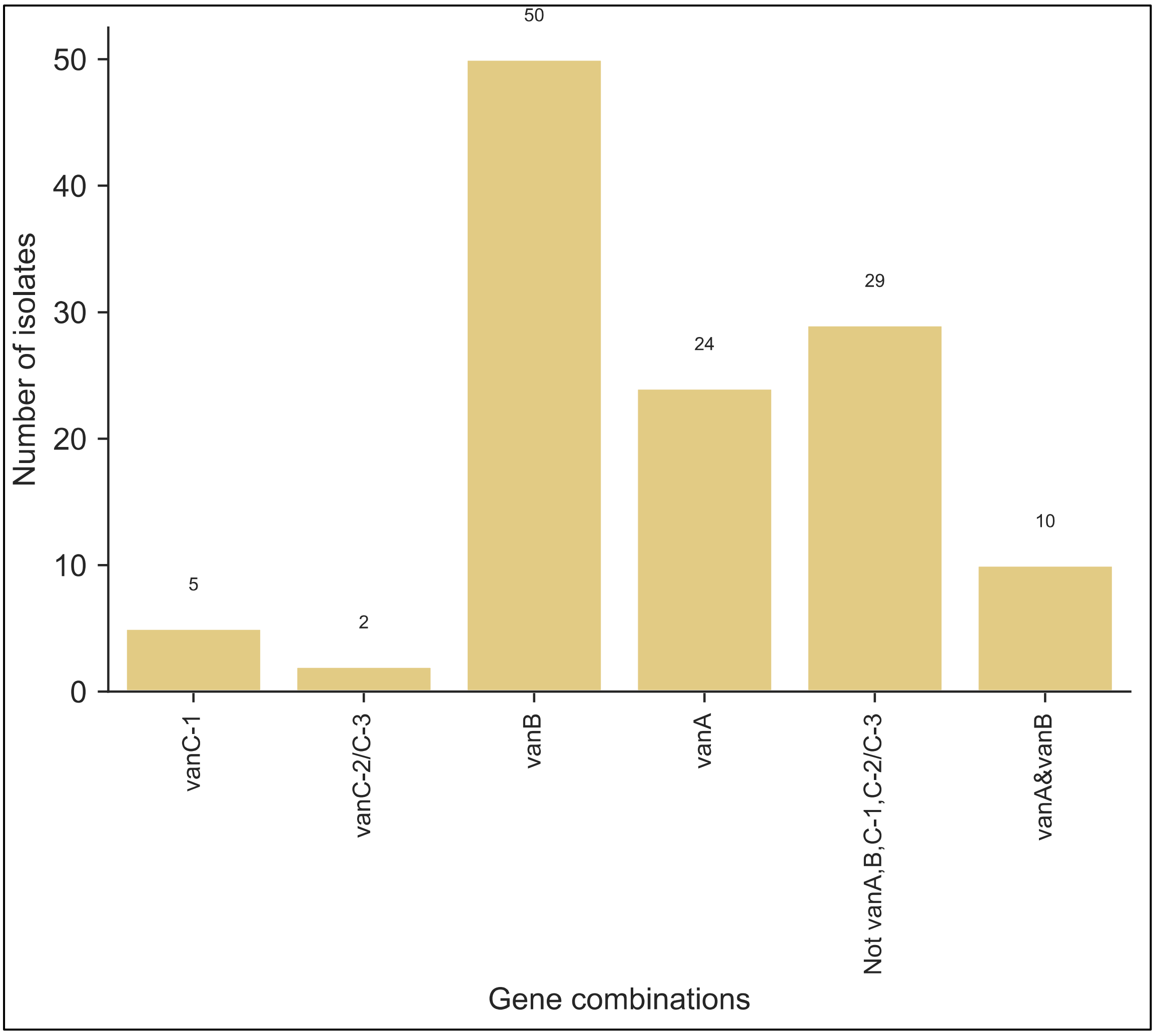

**Figure S7. Resistance alleles used in validation of allele calling**

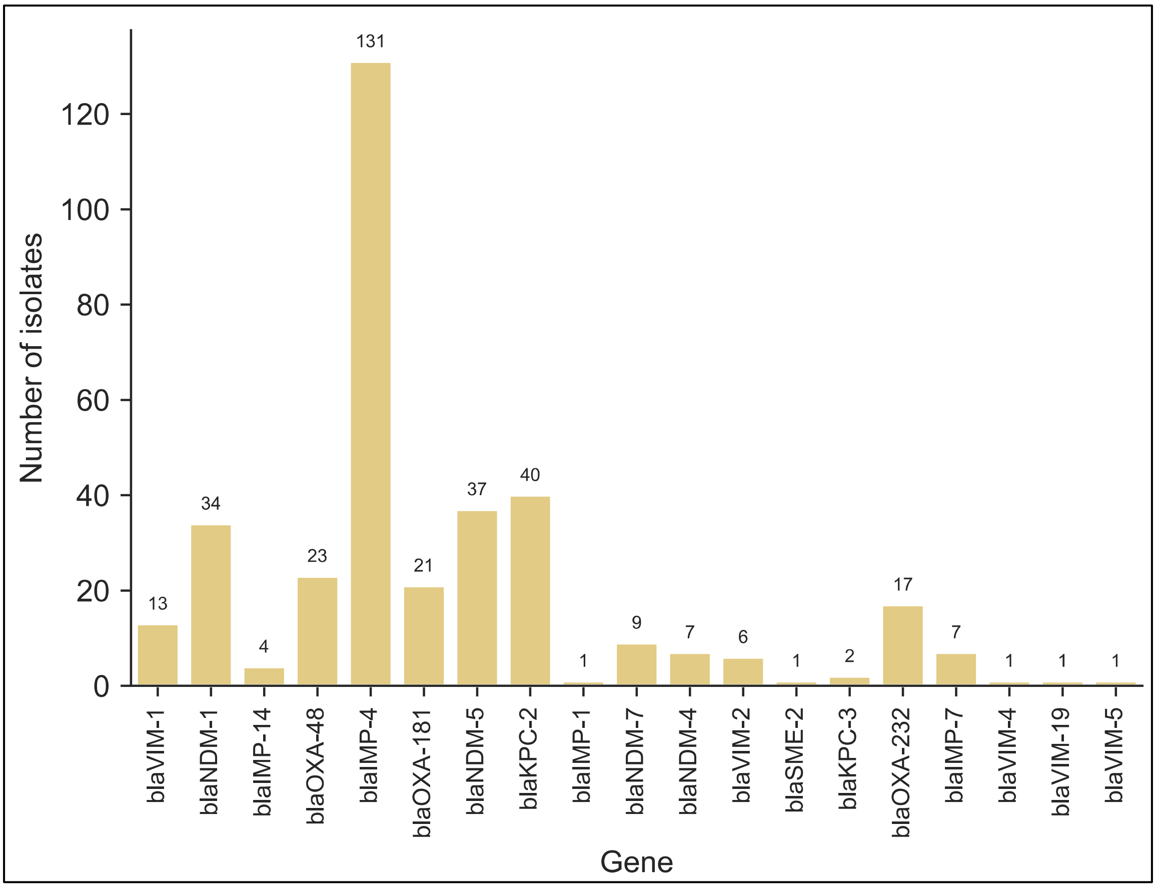

**Figure S8. Genera and species included in validation of resistance allele calling**

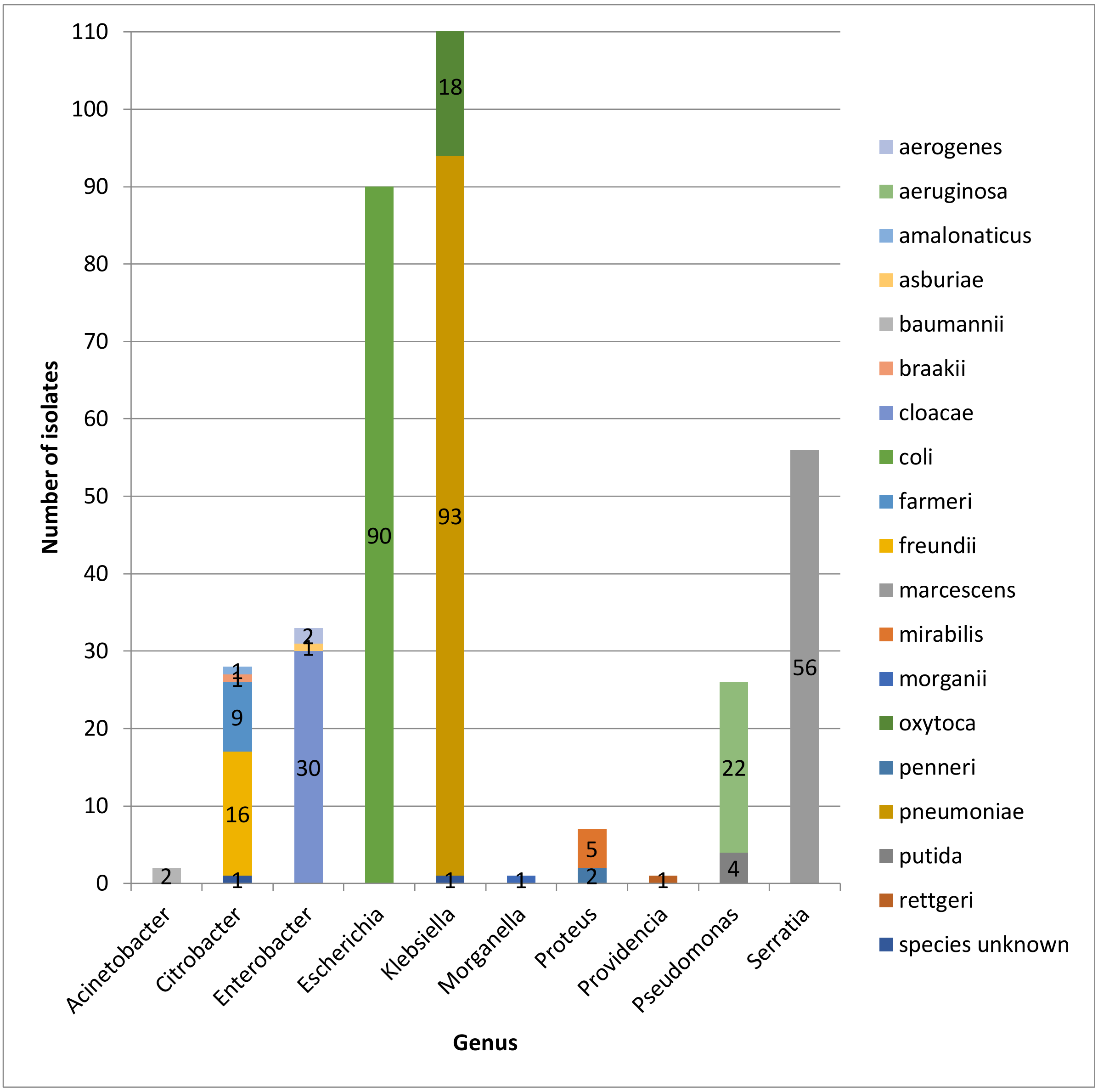

**Figure S9. Genera and species included in validation of resistance allele calling from synthetic dataset**

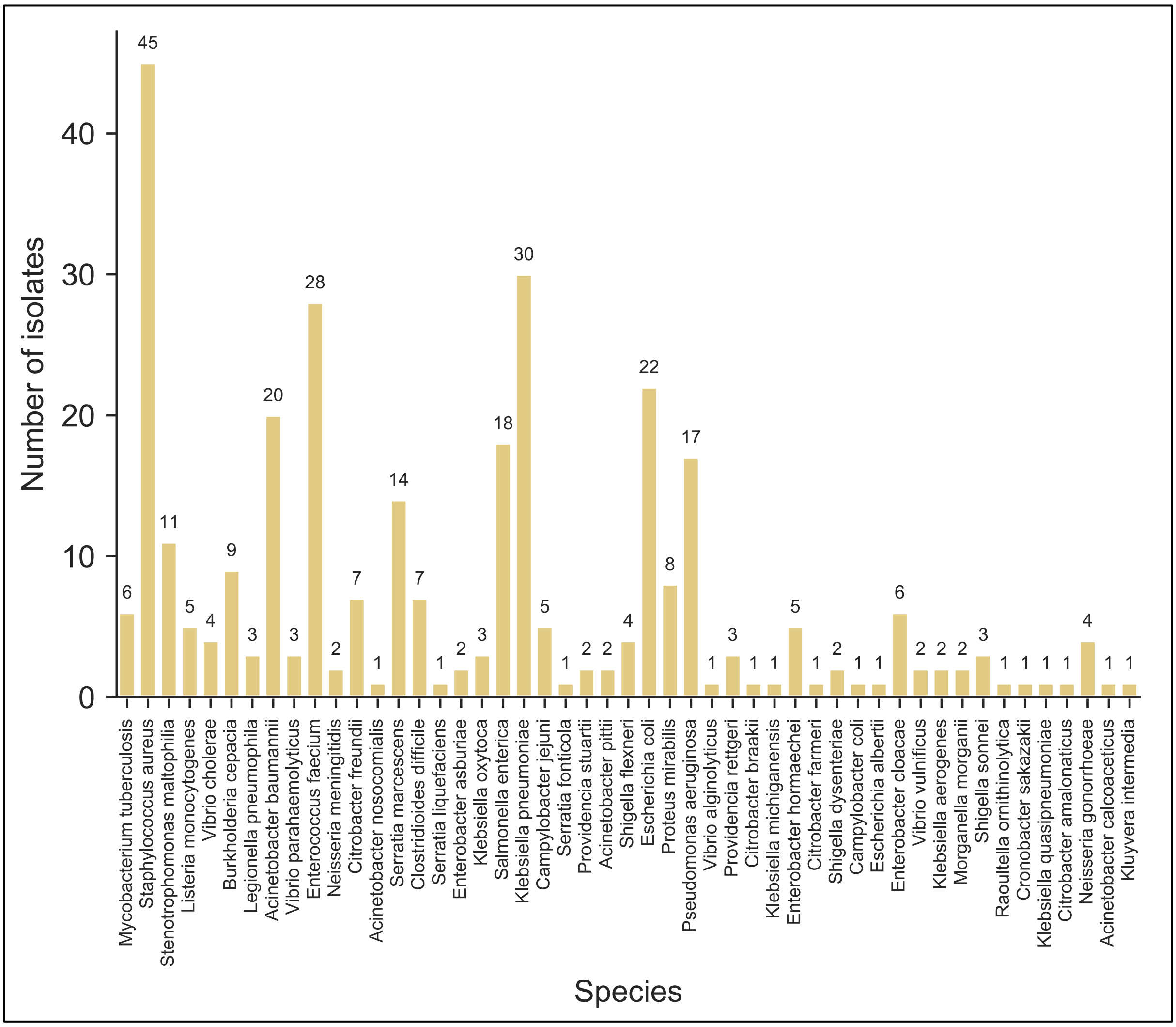

**Figure S10. Drug classes with represented in synthetic dataset for the validation of resistance allele calling**

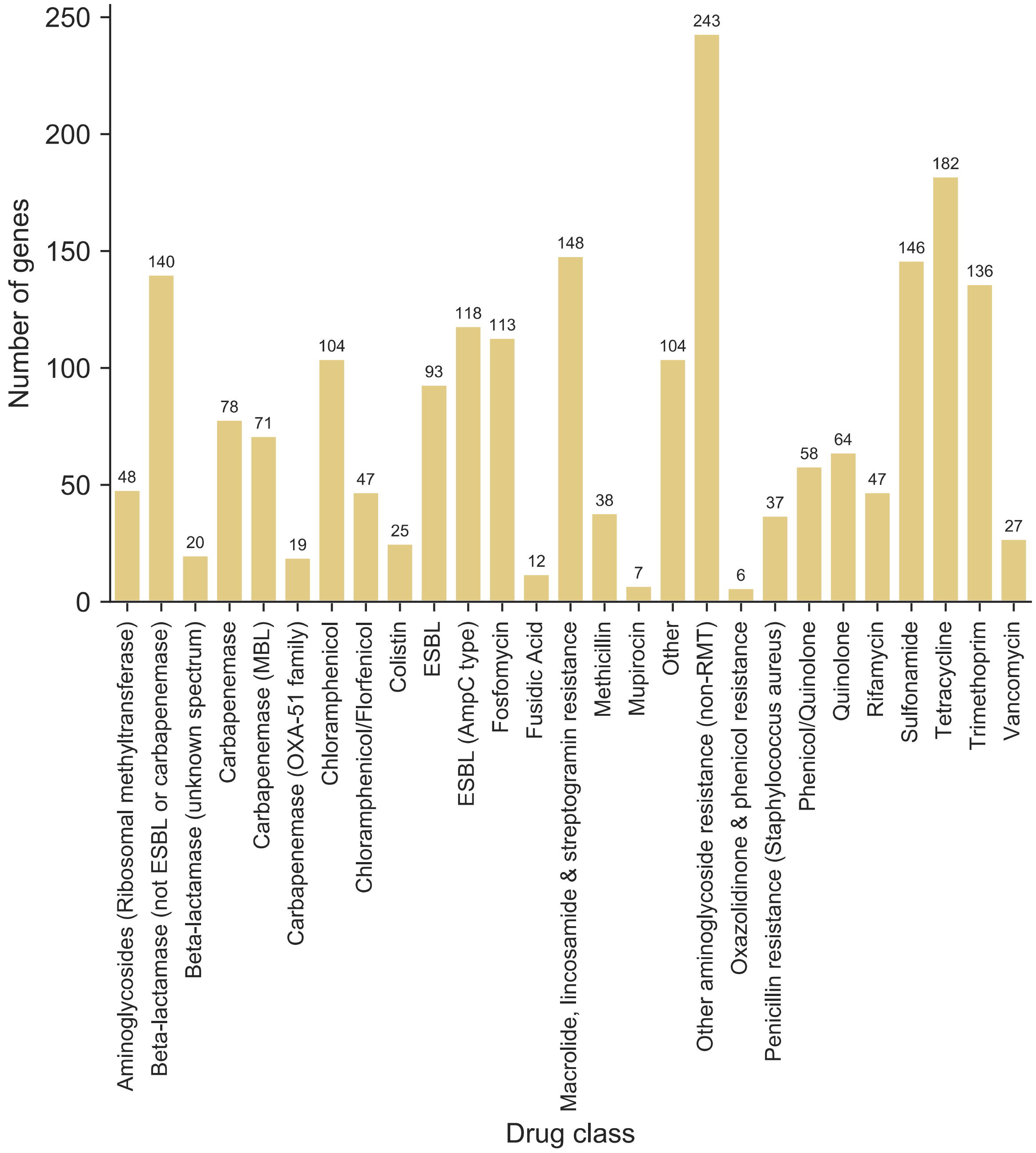
